# Caspase-Driven Microvascular Inflammation and Hypoperfusion in Intravascular Hemolysis: Roles of Leukocyte and Endothelial Activation

**DOI:** 10.1101/2025.03.10.642471

**Authors:** Pamela L. Brito, Lucas F.S. Gushiken, Erica M.F. Gotardo, Vanessa Figueiredo, Flavia C. Leonardo, Fernando F. Costa, Nicola Conran

**Author notes:** **Corresponding author:** Nicola Conran, Hematology Center, University of Campinas, Barao Geraldo, Campinas, São Paulo, Brazil.

## Abstract

Intravascular hemolysis (IVH), a pathological process associated with various conditions, triggers vascular responses; however, the molecular and cellular mechanisms driving this process remain unclear. To explore the role of NLRP3 inflammasome- and caspase-dependent pathways in IVH-induced inflammation, we used *in vivo* models of acute and chronic IVH, alongside heme-stimulation of endothelial cells, thereby isolating this disease mechanism from its etiological causes. Acute IVH induced immediate inflammatory responses in C57BL/6 mice, marked by the release of pro-inflammatory molecules, including IL-1β, within just 15 minutes and NLRP3-dependent caspase activation in circulating leukocytes. Chronic IVH processes in mice elevated liver monocyte-derived macrophage caspase activity and NLRP3 protein expression. In turn, acute IVH impaired cutaneous microvascular blood flow and perfusion, and induced microvascular leukocyte recruitment, which were both caspase-1-dependent. Acute IVH induced time-dependent CD11b-integrin-subunit presentation on leukocytes, while heme stimulation augmented endothelial cell adhesion molecule expression, potentially promoting leukocyte recruitment. This endothelial activation was associated with reactive oxygen species generation, which promoted caspase-1 activation and was key to adhesion molecule upregulation. Our findings highlight a role for inflammasome-/caspase-dependent pathways in hemolytic inflammation, contributing to microvascular leukocyte recruitment and particularly to cutaneous hypoperfusion, a consequence that could facilitate the progression of skin lesions. Targeting caspase-dependent pathways and their downstream effects in disorders that display IVH may offer therapeutic potential for maintaining endothelial integrity, reducing leukocyte activation, and mitigating ischemic injury.

**NEW & NOTEWORTHY:** This study identifies caspase-1 as a driver of the vascular inflammation and hypoperfusion induced by intravascular hemolysis (IVH). Using *in vivo* models and heme-stimulated endothelial cells, we show that hemolysis rapidly induces caspase-1-dependent endothelial-leukocyte recruitment, microvascular dysfunction, and also IL-1β release. Oxidative stress promotes heme-induced endothelial caspase-1 activation and adhesion molecule expression, potentially amplifying vascular dysfunction. These findings provide insight into IVH-driven pathology in hemolytic disorders, including sickle cell disease.

## INTRODUCTION

Hemolysis, or the destruction of red blood cells (RBCs) is a major pathological process that can be classified into two primary types: extravascular hemolysis, occurring predominantly in the spleen and the liver, via macrophage-mediated phagocytosis, and intravascular hemolysis (IVH), where RBC destruction occurs directly within the bloodstream (1). As a critical disease mechanism, hemolysis drives systemic inflammation, upregulates coagulation pathways and induces endothelial dysfunction, hallmarks of several disorders with associated hemolysis, such as sickle cell disease (SCD), paroxysmal nocturnal hemoglobinuria (PNH), malaria and hemolytic transfusion reactions (1–3). While these conditions exhibit distinct etiologies and clinical manifestations, they share common hemolysis-related complications, including pulmonary hypertension, cutaneous leg ulcers and priapism (4–6).

Intravascular hemolysis, in particular, has significant systemic effects, due to the uncontrolled release of Hb, heme and other erythrocyte contents into the circulation. Cell-free Hb binds nitric oxide (NO), impairing vasodilation, while arginine released from RBCs reduces NO bioavailability, contributing to endothelial dysfunction and vasoconstriction (7). Typically, this process is controlled by haptoglobin, hemopexin, and other heme-binding proteins, which scavenge free Hb and heme. However, during sustained hemolysis, these protective systems become saturated leading to several consequences, including complement system activation and rapid inflammatory responses (1, 8–11). In addition, free heme, derived from oxidized Hb, promotes oxidative stress and acts as a danger-associated molecular pattern (DAMP) (12, 13) that, together with other RBC-derived DAMPs, may trigger innate immune responses, via Toll-Like receptors (TLR) signaling and inflammasome activation, resulting in the processing and release of pro-inflammatory cytokines, such as interleukin (IL)-1β and IL-18, and neutrophil extracellular traps (NETs)(14–16). Although such molecular mechanisms implicate a major role of IVH in inducing immune responses, the exact mechanisms by which IVH exacerbates the pathologies of hemolytic diseases remain incompletely understood.

We previously standardized two murine models of hemolysis that emulate moderate IVH. The acute IVH model, induced by a single administration of sterile water, triggers hypertonic RBC lysis and leads to vascular inflammation (17). A similar water-infusion protocol in dogs led to rapid NO depletion and endothelial dysfunction (3). In mice, acute IVH induces inflammatory responses that include microvascular leukocyte recruitment and release of the S100A8 alarmin (17, 18). In contrast, our murine chronic IVH model is induced by repetitive low doses of phenylhydrazine (LDPHZ), causing continuous RBC lysis through membrane peroxidation (19). This model results in mild anemia, as evidenced by reticulocytosis and sustained depletion of haptoglobin and hemopexin, ultimately driving inflammatory processes (19). The inflammatory state that is induced by continuous hemolysis is characterized not only by elevated plasma levels of pro-inflammatory molecules, along with endothelial cell activation, but also by pathological hepatic alterations, including increased hepatic iron deposition, sinusoid congestion and inflammatory cell infiltration (19). These models, which mimic moderate hemolytic processes, have proven to be valuable and effective tools to investigate the mechanism of IVH-induced inflammation in hemolytic disorders, independently of the cause of this hemolysis.

Inflammation plays a pivotal role in driving vascular disturbances through a cascade of processes that lead to leukocyte recruitment, endothelial activation, and microvascular occlusion (20). Pro-inflammatory molecules, such as TNF-ɑ and heme, can enhance the expression and activation of adhesion molecules on endothelial cells and leukocytes (21–23), promoting leukocyte adhesion to the vascular wall (24). This response is further amplified by leukocyte mobilization, the release of additional pro-inflammatory cytokines and the generation of reactive oxygen species (ROS) by activated leukocytes (19, 25, 26). Together with NO depletion, generating impaired vasodilation and platelet activation, these inflammatory processes exacerbate vascular dysfunction (27, 28). As such, IVH-driven inflammation and hemodynamic changes may play a critical role in the pathology of hemolytic disorders, where their interplay may be central to vascular dysfunction and microvascular occlusion.

One key mechanism that contributes to innate immune responses is inflammasome activation (29). The multiprotein intracellular inflammasome complexes assemble in immune cells in response to a danger signals, leading to the activation of caspase-1, a protease that cleaves pro-IL-1β and pro-IL-18 into their active forms (30). Active caspase-1 also induces gasdermin-mediated pyroptosis, a form of programmed cell death that amplifies intracellular content and cytokine release (31). Subsequently, IL-1β stimulates endothelial adhesion molecule expression and enhances the production of inflammatory mediators such as IL-6, IL-8 and monocyte chemoattractant protein-1 (MCP-1), thereby contributing to leukocyte recruitment (32, 33). Additionally, NLRP3 inflammasome activation has been reported in non-immune cell types including the endothelial cells themselves and platelets (34, 35). Hemolytic products, such as heme and oxidized Hb, are potent DAMPs that are known to trigger NLRP3 inflammasome activation in lipopolysaccharide (LPS)-primed macrophages and endothelial cells, leading to IL-1β maturation and release (29, 36).

We hypothesized that inflammasome activation plays a key role in the rapid multicellular vascular inflammatory responses to acute IVH, contributing to vascular inflammation and potentially to hemodynamic alterations. Over time, persistent hemolytic processes may drive some of the long-term vascular complications associated with chronic hemolytic diseases. To address this, we employed our animal models of hemolysis to replicate moderate acute and chronic hemolysis without interference from underlying pathologies, allowing us to directly assess the impact of the hemolytic process in vascular dysfunction. Additionally, we used *in vitro* assays to explore mechanisms of heme-stimulated macrovascular and microvascular endothelial activation.

## METHODS

### Animals and Ethics Information

C57BL/6J and transgenic Townes mice, maintained at the animal facility (CEMIB) of the University of Campinas (UNICAMP), were used aged 4 to 5 months for the study. The animals were housed in microisolators at until euthanasia, with food and water provided *ad libitum*. All procedures were conducted in compliance with national guidelines for animal experimentation and the study was approved by the Ethics Committee on Animal Use of UNICAMP under protocols 5144-1/2019 and 5144-1(A)/2022. All efforts were made to minimize animal suffering and to use the minimum number of animals necessary to obtain reliable and replicable results. Animals were euthanized at the end of final procedures by deep anesthesia (300 mg/kg ketamine and 30 mg/kg xylazine, *i.p*.). This study exclusively used male mice due to the necessity to perform intravital microscopy of a surgical preparation of the cremaster muscle.

### Acute and Chronic Intravascular Hemolysis Induction

Induction of hemolytic processes was performed as previously described (17, 37). For the induction of acute IVH, isogenic C57BL6/J mice received a single hemolytic stimulus via the *i.v.* administration of 150 µL of sterile water. As controls, mice of the same strain received an injection of sterile saline solution (0.9% NaCl). Experimental groups were defined based on the duration of hemolysis, with sample collection performed at 15 minutes, 1 hour, 3 hours and 6 hours after water/saline administration. For the induction of chronic hemolysis, isogenic C57BL6/J mice received intraperitoneal (*i.p*.) injections of a low dose of phenylhydrazine (10 mg/kg; PHZ, Sigma-Aldrich) twice weekly for four weeks, totaling 8 administrations. Similarly, control animals without hemolysis received *i.p.* sterile saline solution (0.9% NaCl) administrations. Mice from all groups were euthanized 48 hours after the last PHZ administration for sample collection.

### Animal Treatment Protocols

Prior to the induction of acute IVH or saline administration, mice were treated for 1 hour with the NLRP3 inflammasome inhibitor, MCC950 (50 mg/kg, *i.p*, Sigma-Aldrich), while control groups received sterile phosphate-buffered saline (PBS). For caspase-1 inhibition, mice were treated with AC-YVAD-cmk (8 mg/kg, Sigma-Aldrich) or 0.5% Dimethyl Sulfoxide vehicle (DMSO, Sigma-Aldrich) in sterile PBS, via intraperitoneal injection.

### Laser Doppler Flowmetry

Cutaneous microvascular blood velocity and perfusion were assessed in the depilated pelvic region of anesthetized mice (80 mg/kg ketamine/10 mg/kg xylazine, *i.p*.) using the PeriFlux 6000 system (Perimed Instruments, Järfälla, Sweden). Blood cell concentration (CMBC), velocity, and perfusion were assessed/calculated for 10 minutes at a depth of 0.3 to 0.5mm, using a Ø1.0 mm diameter probe with a 780 nm laser (38). Baseline measurements were taken 30 minutes before hemolysis induction or treatments, and at 15 minutes or 1 hour after acute hemolysis induction or saline administration. Absolute values were normalized to each animal’s baseline condition.

### Intravital Microscopy

After surgical externalization of the cremaster muscle (38) in anesthetized mice (ketamine/xylazine*, i.p*.), animals were placed under a microscope (Zeiss Imager M.2, Carl Zeiss Microscopy, Jena, Germany) for visualization of the microvasculature. Images from 6 to 8 venules per animal, with diameters ranging between 25 and 35 µm, were acquired using an Axiocam 506 color camera (Carl Zeiss Microscopy) and the Zen 2 Pro software (Carl Zeiss Microscopy). Blood flow images (40 frames per second) were captured using a water-immersive 63X objective under brightfield illumination for a maximum of 20 minutes. Video segments of 1-minute duration were exported for quantitative analysis of rolling, adherent, and extravasated leukocytes (19).

### Isolation of Hepatic Non-Parenchymal Cells

For non-parenchymal cell (NPC) isolation, mouse livers were perfused *in situ* with perfusion solution (Liver Perfusion Medium, 37°C, Gibco™, MA USA), after which the liver was gently dissected and digested with buffer containing collagenase (Liver Digest Medium, Gibco) at 37°C for 40 minutes, with occasional agitation. After filtering (100-μm nylon cell strainer), homogenates were washed three times (PBS, 30 × g for 4 minutes at 25°C) for hepatocyte and debris sedimentation. The supernatant was then collected and subjected to two consecutive centrifugations (300 × g, 10 minutes, 22°C) to sediment non-parenchymal cells (NPCs), including leukocytes and macrophages (39). The collected NPC fraction was carefully resuspended in PBS, and centrifuged at 1500 × g for 20 minutes at 25°C. The resulting pellet was resuspended in sterile RPMI medium.

### Endothelial Cell Culture and Treatments

Human Umbilical Vein Endothelial Cells (HUVECs) and Human Microvascular Endothelial Cells (HMEC-1) were acquired from the American Type Culture Collection (ATCC®, VA, USA). HUVECs were cultured in modified Ham’s F-12 Kaighn’s medium (F12K, Gibco) with 10% fetal bovine serum (FBS, Invitrogen, MA, USA), endothelial growth factor (30 ng/mL, Calbiochem, CA, USA), sodium heparin (10 U/mL, Cristália, São Paulo, Brazil), and antibiotics (streptomycin and penicillin). HMEC-1 were cultured in MCDB 131 medium (Gibco) with 10% FBS, 10 ng/mL Epidermal Growth Factor (EGF, Thermofisher, MA, USA), 1 µg/mL hydrocortisone (Sigma-Aldrich), 10 mM L-Glutamine (Gibco), and antibiotics. Both cell types were maintained in a humidified atmosphere at 37°C with 5% CO₂, and used at 80-90% confluence, between 3 and 6 passages. For experiments, HUVEC or HMEC-1 were seeded at 2 × 10⁵ cells/mL per 9.6 cm² well in complete F12K or MCDB 131 medium supplemented with 2% FBS, for 24 hours at 37 °C, 5% CO₂. Cells were stimulated, or not, with heme (50 µM; Frontier Scientific, UT, USA) for 3 hours at 37°C, 5% CO₂. For caspase-1 inhibition, cells were pretreated for 1 hour with YVAD-cmk (50 µM, Sigma-Aldrich) or 0.25 % dimethyl sulfoxide vehicle (DMSO, Sigma-Aldrich) before heme stimulation for an additional 3 hours. For antioxidant treatment, cells were incubated with 1 mM α-tocopherol (DL-all-rac-α-tocopherol, Vitamin E, Sigma-Aldrich), or 2% ultrapure ethanol vehicle, for 30 minutes prior to stimulation with heme.

### Flow Cytometry: Murine Cells

Cellular caspase-1 activation and cell phenotyping was determined in murine cells using flow cytometry. Fifty μL of EDTA-coagulated whole blood (for neutrophil and monocyte analysis) and NPC samples (1 × 10⁶ cells/mL) were pre-incubated with a blocking solution (Mouse BD Fc Block purified anti-mouse CD16/CD32 – 1:50, BD Pharmingen) for 15 minutes at room temperature before incubating with the intracellular staining probe, FAM-YVAD-FMK (FAM-FLICA Caspase-1, ImmunoChemistry Technologies, Bloomington, MN, USA), and phenotyping antibodies (37 °C, 30 minutes). After incubation, red blood cells were lysed using BD Pharm Lyse Buffer (BD Bioscience; 15 minutes at room temperature). Immediately after lysis, cells were washed (Apoptosis Wash Buffer; FAM-FLICA Caspase-1 (YVAD) Assay Kit) and resuspended in PBS containing fixative solution (FAM-FLICA Caspase-1 (YVAD) Assay Kit) for acquisition on a FACSCalibur flow cytometer (BD Biosciences, CA, USA), with data analyzed by FlowJo software v.10.5.3. Viable neutrophils (Ly6G^high^ CD11b⁺), monocytes (Ly6C^high^ CD43⁺), and liver macrophages (F4/80⁺ CD11b⁺) were identified after gating based on SSC/FSC parameters. Caspase-1 activation was expressed as the percentage of phenotyped FLICA⁺ cells (5x10^4^ events). All antibodies used in flow cytometry are listed in Supplementary Table 1.

### Flow Cytometry: Endothelial Cells

For analysis of adhesion molecules and caspase-1 activity, HUVEC and HMEC-1 cells (1 × 10^5^ cells/mL) were fixed with 1% paraformaldehyde, then dissociated with 0.25% Trypsin/0.53 mM EDTA solution (37°C) prior to antibody and FAM-FLICA Caspase-1 probe labelling for 30 min, 37°C (see Supplementary Table 1). For ROS measurement, unfixed cells were collected, washed with PBS, and immediately incubated with the 2′,7′-Dichlorodihydrofluorescein diacetate probe (DCFH-DA, Sigma-Aldrich; 10 µM, 30 minutes, 37 °C). Event acquisition with a FACSCalibur (2 × 10⁴ events), and data were analyzed using FlowJo software v.10.5.3. The gating strategy used SSC/FSC (side scatter/forward scatter) parameters to identify viable cells and the population of interest.

### Immunoassay Protein Measurements

Murine plasma samples were used for enzyme-linked immunosorbent assays (ELISA) to quantify IL-1β (mouse IL-1β/IL-1F2 [high sensitivity] ELISA Kit, R&D), IL-18 (mouse IL-18 ELISA Kit, MBL), IL-6 (mouse IL-6, Abcam), TNF-α (Mouse TNF-alpha Quantikine [high sensitivity] ELISA Kit, R&D), S100A8 protein (DuoSet Mouse S100A8, R&D), haptoglobin, and hemopexin (Abcam), following the manufacturers’ instructions. Total liver protein extracts were used to quantify hepatic IL-1β using the commercial mouse IL-1β SimpleStep ELISA kit (Abcam), according to the manufacturer’s instructions.

### Hemoglobin and Heme Measurement

Plasma hemoglobin was determined according to the method of Fairbanks et al. (40); hemoglobin levels are expressed as µM heme bound to hemoglobin. Total cell-free plasma heme was measured using a commercial kit (QuantiChrom™ Heme Assay Kit BioAssay Systems, San Francisco, CA), according to The Manufacturer’s Instructions.

### Quantitative Real Time PCR (qPCR)

mRNA from HUVECs was isolated using the RNeasy Micro Kit (Qiagen, Germany), according to the manufacturer’s instructions. cDNA synthesis was performed using the RevertAid H Minus First Strand cDNA Synthesis Kit (Thermo Fisher Scientific) and cDNA concentration measured using a NanoDrop spectrophotometer (Thermo Fisher Scientific). All qPCR samples were run in duplicate on the same plate in a total volume of 12 µL, containing 10 ng of cDNA, 6 µL of SYBR Green Master Mix PCR (Applied Biosystems, CA, USA), and oligonucleotide primers for the genes of interest (*CASP1*, *NLRP3* and *PYCARD*, designed by Primer-Express, Applied Biosystems, and synthesized by IDT, Iowa, USA). For oligonucleotide primers sequences and concentrations, see Supplementary Table 2. The reaction was performed using the 7500 Fast Dx Real-Time PCR instrument and software (Applied Biosystems). To confirm the accuracy and reproducibility of the qPCR, intra-assay precision was calculated using the equation E(-1/slope), and a dissociation curve analysis was performed at the end of each run to detect nonspecific amplifications. Results are expressed as arbitrary units (A.U) of gene expression and were normalized to β-actin and GAPDH expression, calculated using the geNorm program (41).

### Protein Extraction and Immunoblotting

Liver fragments were homogenized in lysis buffer, containing a protease inhibitor cocktail. Total protein was determined using the Bradford colorimetric method. Fifty μg of protein extract were loaded onto a 4-20% polyacrylamide gel (SDS-PAGE – Premade MiniProtean gel, Bio-Rad) for western blotting. The following primary antibodies (diluted 1:10) were used: rat anti-murine NLRP3 monoclonal antibody (MAB7578; R&D), rabbit anti-IL-1β polyclonal antibody (ab9722; Abcam), rabbit anti-caspase-1 polyclonal antibody (ab138483; Abcam), and mouse anti-GAPDH monoclonal antibody (sc-365062; Santa Cruz). Secondary antibodies (diluted 1:1000) were anti-rat IgG-HRP-conjugated (sc-2006; Santa Cruz), anti-rabbit IgG-HRP-conjugated (sc-2357; Santa Cruz) and anti-mouse IgG-HRP-conjugated (sc-358914; Santa Cruz). Blots were developed using the Pierce ECL chemiluminescence detection kit (Thermo Scientific, USA). Images were visualized and captured using a photo documentation system (UVISoft-UVIBand, UVITEC Cambridge ALLIANCE-6.7); band intensity values were normalized to the optical densities of GAPDH bands, and analyzed with ImageJ software.

### Statistical Analysis

Data are expressed as means and standard error of N samples. For comparisons between two groups, the Mann-Whitney test was used. For comparisons between three groups or more, data normality was assessed, and either a One-Way Analysis of Variance (ANOVA) test followed by Dunn’s multiple-comparison test (for non-parametric samples) or Holm’s-Sidak’s multiple comparison test (for parametric samples) was employed using the GraphPad InStat software. A probability (P value) of 5% or less was considered significant.

## RESULTS

### A Single Acute Hemolytic Event Induces a Rapid Pro-Inflammatory Effector Response in Mice

An acute intravascular hemolytic event was induced in C57BL/6 mice by injecting them (i.v.) with sterile water, according to a previously established protocol (42). A control group received a single administration of sterile saline. Blood samples were collected at 15 min, 1 hour and 3 hours later (Figure 1). As previously described (42), this protocol rapidly increases plasma cell-free hemoglobin and total plasma cell-free heme, confirming the occurrence of IVH within 15min (Figure 1A and B). Plasma haptoglobin was also rapidly depleted within 1 hour (Figure 1C). In association with these observations, acute IVH triggered prompt inflammatory protein release, with plasma levels of TNF-α cytokine increasing at 3 hours (Figure 1D). Furthermore, circulating levels of the leukocyte-derived S100A8, (Figure 1E) an alarmin that can participate in inflammasome activation, increased within just 15 minutes of the hemolytic stimulus. Inflammasome activation is necessary for the maturation of the IL-1β and IL-18 cytokines; consistent with the hypothesis of inflammasome-mediated cytokine processing, plasma levels of IL-1β augmented within just 15 min of IVH induction (Figure 1F), while IL-18 release was slower, occurring at approximately 6 hours post-hemolysis (Figure 1H). Additionally, liver IL-1β protein content presented a significant increase within the first hour post-hemolysis (Figure 1G).

**Figure 1.**
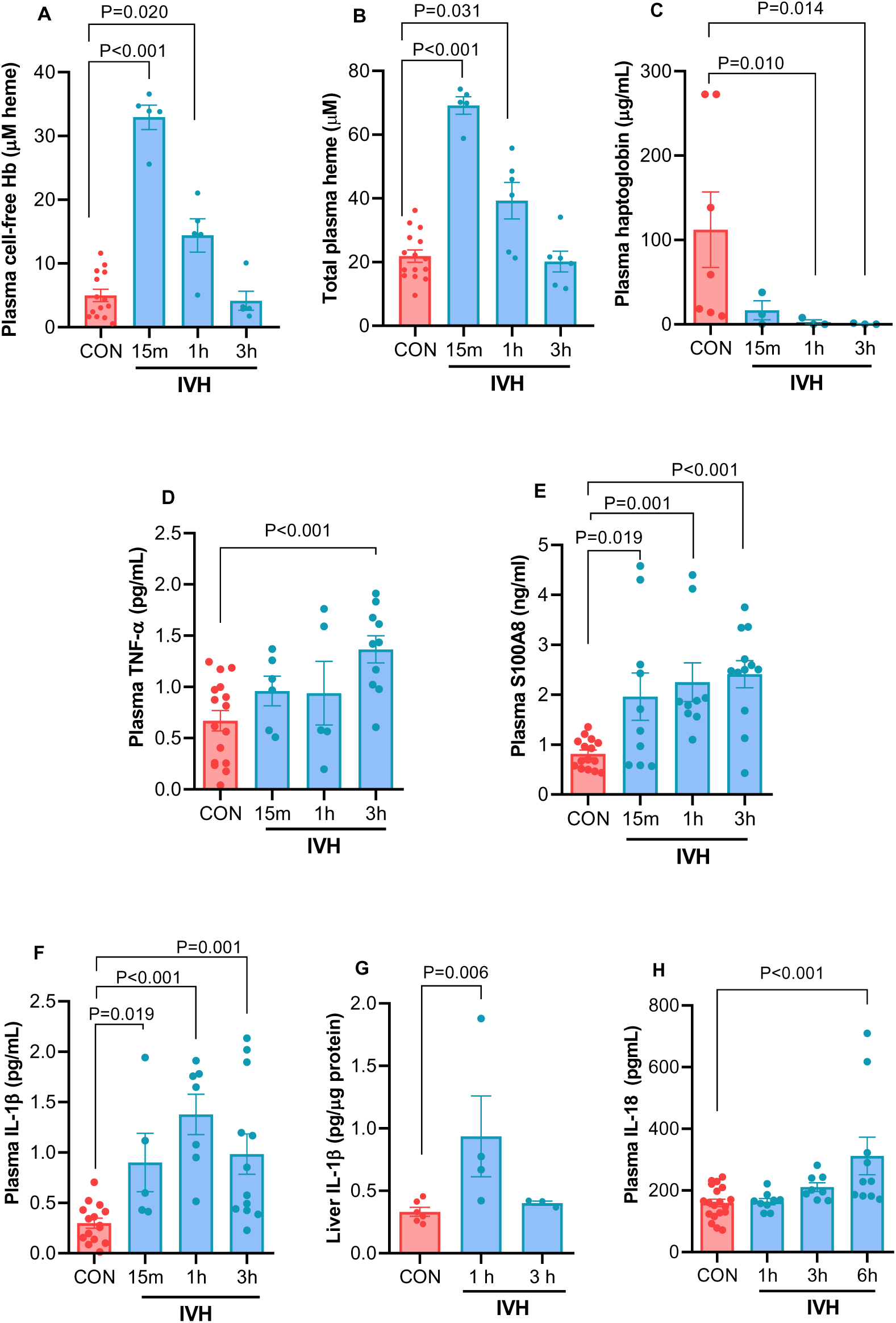
Acute intravascular hemolysis induces rapid inflammatory responses in C57BL/6 Mice. Mice received single intravenous administrations of sterile 0.9% NaCl (CON, total n = 6-19) or sterile water (IVH, n = 3-12) and blood samples were collected at 15 minutes, 1 hour, 3 hours, and 6 hours after administrations for analysis. Saline control samples from different timepoints were pooled into a single group for analysis, while IVH samples were analyzed separately for each timepoint. Plasma cell-free hemoglobin (A) and cell-free heme (B) were measured using the Fairbanks and Quantichrom™ colorimetric assays, respectively, and expressed as μM heme. Plasma Haptoglobin (C), TNF-α (D), S100A8 (E), and IL-1β (F), as well as liver IL-1β (G) and plasma IL-18 (H) were determined by ELISA.

### Acute *in vivo* Intravascular Hemolysis induces NLRP3-Dependent Caspase-1 Activation in Murine Leukocytes

Having observed rapid hemolysis-driven inflammasome-derived cytokine release, we compared caspase activity in the neutrophils and monocytes of mice that had received a single intravenous administration of either sterile 0.9% NaCl (control mice) or sterile water (IVH) to induce IVH (Figure 2). C57BL/6 mice were also pretreated (1 hour before hemolysis) intraperitoneally with either the irreversible caspase-1 inhibitor, YVAD, or the selective NLRP3-inflammmasome inhibitor, MCC950, or their vehicles (*i.p.* DMSO or sterile PBS, respectively). Caspase activity was determined in immunophenotyped neutrophils and monocytes at 1 hour post hemolysis (or saline infusion) using the FAM-YVAD-FLICA-FITC probe, with primary specificity for caspase-1, and flow cytometry.

**Figure 2.**
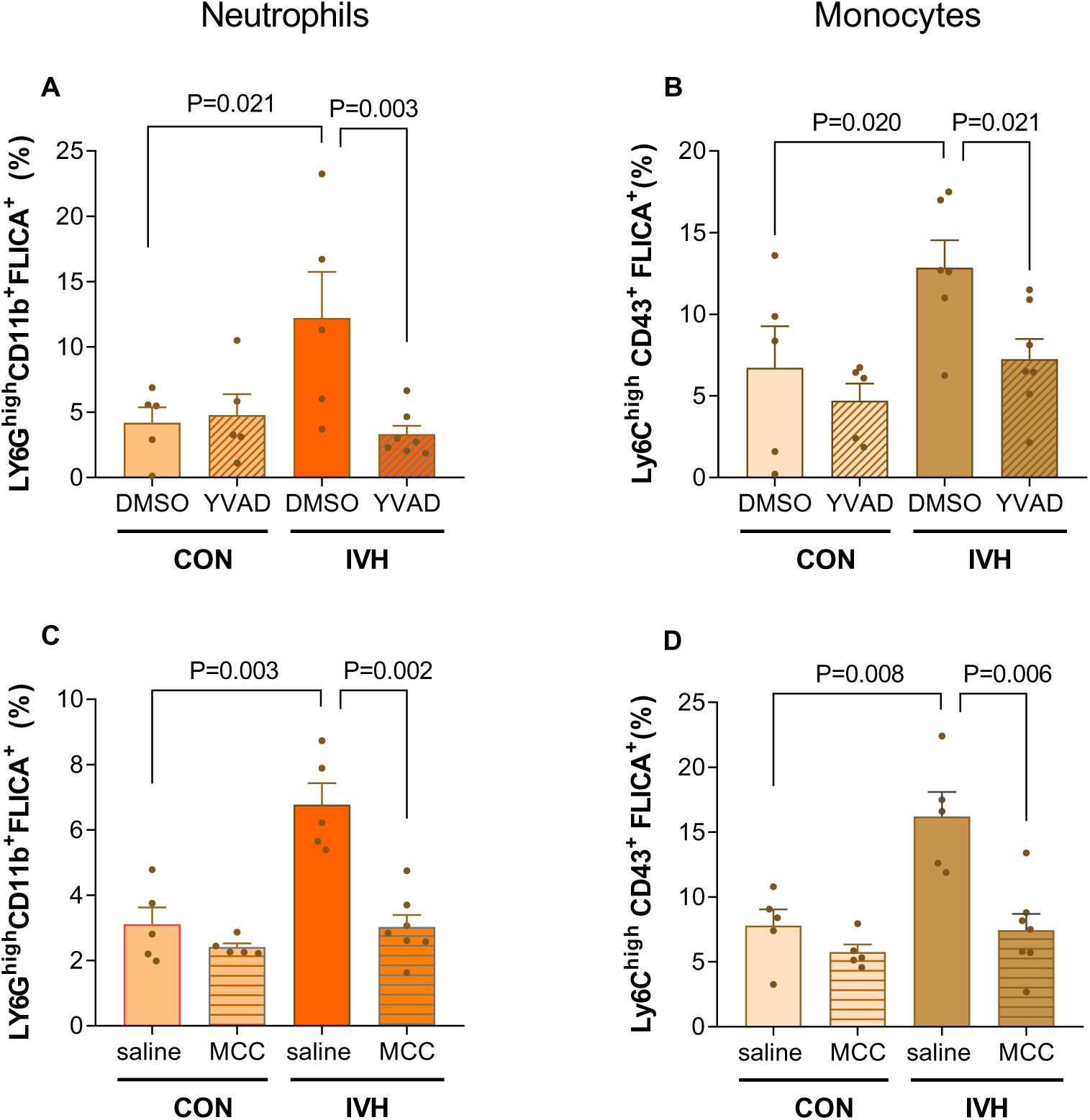
Acute intravascular hemolysis induces NLRP3-dependent caspase-1 activation in murine leukocytes. C57BL/6 mice were pretreated (*i.p*.) with YVAD (caspase-1 inhibitor, 8 mg/kg, A, B) or MCC950 (MCC; NLRP3 Inflammasome Inhibitor; 50 mg/kg, B, D) or with DMSO vehicle (0.5% in sterile PBS, vehicle for YVAD), or with sterile PBS vehicle. One hour later, animals received single intravenous administrations of sterile 0.9% NaCl (CON, n = 5) or sterile water (IVH, n = 5-7) and blood samples were collected 1 hour later for detection of active caspase-1 using FAM-YVAD-FLICA-FITC and flow cytometry. Neutrophils were identified using anti-Ly6G-APC and anti-CD11b-PE antibodies (A, C), while monocytes were identified with anti-Ly6C-APC and anti-CD43-PE antibodies (B, D).

Acute IVH significantly increased caspase activity in both the neutrophils (Ly6G^high^CD11b^+^, Figure 2A, C) and monocytes (Ly6C^high^CD11b^+^, Figure 2B, D) of mice. Importantly, this effect was prevented by both ac-YVAD-CMK (Figure 2A, B) and by MCC950 (Figure 2C, D). While the FLICA probe can detect the activity of other caspases (albeit with lower affinity), the combined inhibition by YVAD and MCC950 strongly suggests that acute IVH induces NLRP3-dependent and, therefore, presumably caspase-1 dependent activity in murine leukocytes.

### Rapid Microvascular Leukocyte Recruitment in Response to Acute Intravascular Hemolysis is Partially Mediated by Caspase-1 Activity

The acute inflammatory response that is triggered by IVH rapidly induces leukocyte recruitment to the microvascular wall at 15-35 minutes post-hemolysis, significantly increasing the rolling and adhesion of these cells at the vascular wall and also increasing extravasation (Figure 3D-F), compared to saline-control mice. When mice were pretreated (1 hour, *i.p*.) with the caspase-1 inhibitor, YVAD, both the rolling and adhesion of leukocytes in response to hemolysis were abrogated (Figure 3A, B), although leukocyte extravasation was not significantly modulated (Figure 3C). In contrast, the decrease in hemolysis-induced leukocyte rolling and adhesion observed following pre-administration of the NLRP3 inhibitor, MCC950, was not statistically significant (Figure 3D-F).

**Figure 3.**
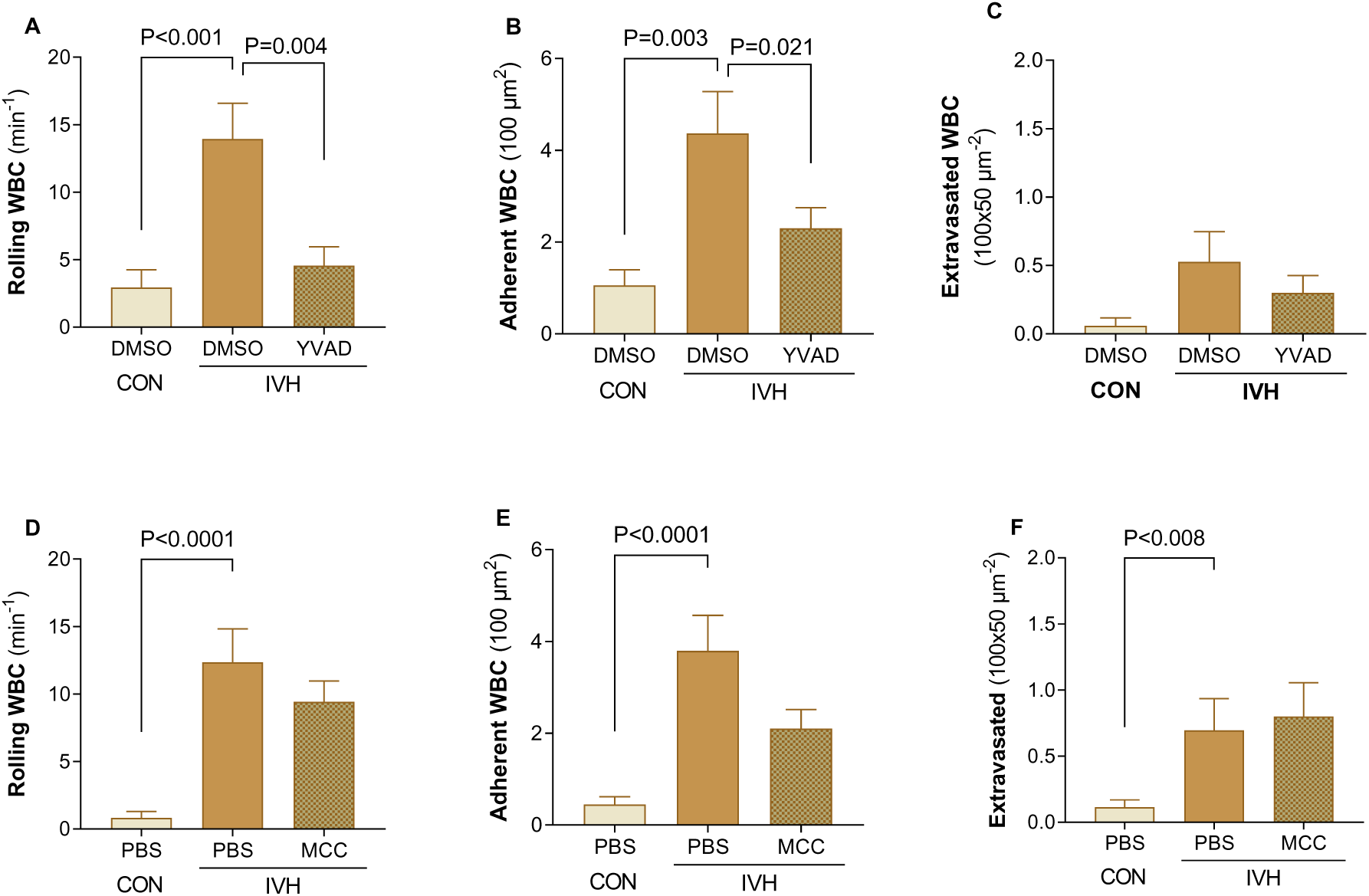
Effects of caspase-1 and NLRP3 inhibition on leukocyte microvascular recruitment in mice subjected to acute intravascular hemolysis. C57BL/6 mice were pretreated (*i.p*., 1 hour) with YVAD (8 mg/kg; A, B, C) or MCC950 (MCC; 50 mg/kg; D, E, F) or with DMSO vehicle (0.5% in sterile PBS, vehicle for YVAD), or with sterile PBS (vehicle for MCC950). Intravital microscopy was performed on the cremaster muscle at 15-35 minutes after intravenous administration of sterile 0.9% NaCl (CON) or after the hemolytic stimulus (IVH). Results are expressed as means ± SEM from a total of 6-8 venules analyzed per animal and n=4 animals in each group. Links for representative videos of each experimental group are provided in Supplementary Material.

In association with these findings, we found no significant effect of IVH on the expression of leukocyte CD11b at 15 minutes post induction; however, by 1 hour CD11b expression on leukocytes was significantly augmented. Flow cytometry demonstrated that the percentage of CD11b^high^ granulocytes and monocytes of mice that received saline was 63.2±3.4 % (n=35), while this percentage was 62.6±3.6 % (n=13) at 15 min and 77.0±3.2 % (n=22, *p*<0.008 compared to saline) at 1 hour post-IVH induction.

### Acute *in vivo* Intravascular Hemolysis Significantly Impairs Microcirculatory Blood Perfusion in an NLRP3- and Caspase-1-Dependent Manner

Since we observed an effect of acute IVH upon microvascular leukocyte recruitment, we investigated the impact of this event on microvascular blood flow and perfusion. Laser Doppler flowmetry was used to monitor the cutaneous blood flow and perfusion in C57BL/6 mice at 15-35 min minutes after the induction, or not, of acute IVH (Figure 4). Mice were also pretreated 1 hour prior to hemolysis induction with intraperitoneal YVAD or MCC950, and vehicle control groups were given DMSO or sterile PBS, respectively.

**Figure 4.**
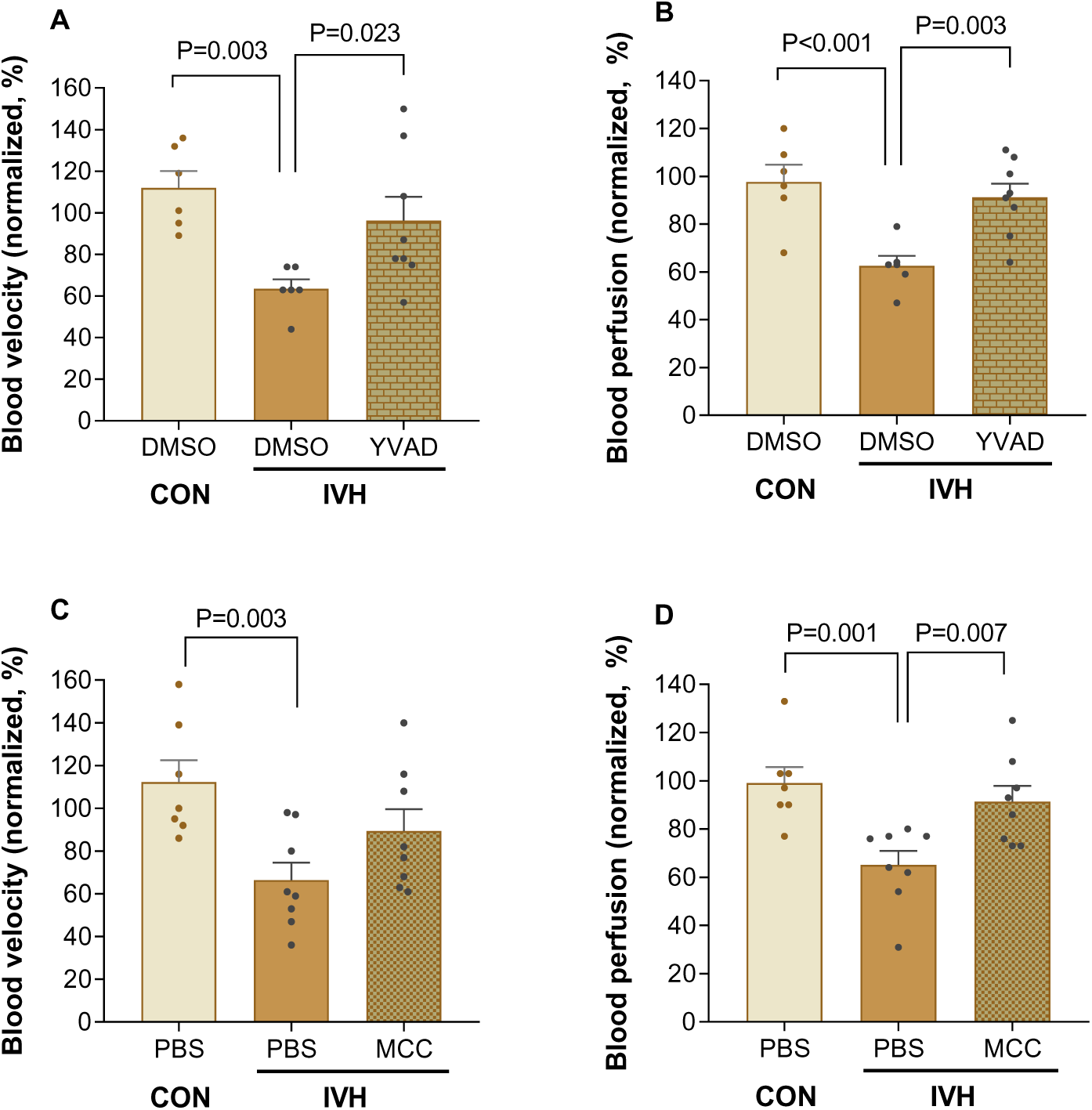
Acute intravascular hemolysis induces NLPR3- and caspase-dependent impairment of cutaneous blood microcirculation hemodynamics in C57BL/6 mice. C57BL/6 mice were pretreated (*i.p*., 1 hour) with YVAD (8 mg/kg; A, B) or MCC950 (MCC; 50 mg/kg; B, D) or with DMSO vehicle (0.5% in sterile PBS, vehicle for YVAD), or with sterile PBS (vehicle for MCC950). Non-invasive Laser Doppler flowmetry was used to measure peripheral microcirculatory blood flow (A, C) and perfusion (B, D) in the skin of the hind limb of anesthetized mice, at 15-35 minutes after intravenous administration of sterile 0.9% NaCl (CON, n = 6-7) or sterile water (IHV, n = 6-9). Data (in arbitrary units) are normalized relative to baseline unstimulated measurements taken from the same mice, at 30 minutes before treatments.

The induction of acute IVH in mice induced a very rapid and significant hypoperfusion in the skin of mice, decreasing both blood velocity (Figure 4A, C) and blood perfusion (Figure 4B, D). Importantly, both caspase-1 (Figure 4A, B) and NLRP3 (Figure 4C, D) inhibition were able to partially or fully restore the cutaneous microvascular blood flow and perfusion in mice subjected to acute IVH, implying a role for the inflammatory response and NLRP3-inflammasome activation in the ischemic response observed.

### Chronic Intravascular Hemolysis in Mice Induces an Hepatic Inflammatory Response and Inflammasome-Derived Cytokine Release

Using a previously-established model (19), we induced long-term continuous IVH in C57BL/6 mice by treating them for 28 days with low-dose phenylhydrazine (10 mg/kg *i.p*., 2x per week). Control mice received *i.p.* saline at the same administration frequency. At 48 h after the last phenylhydrazine dose, mice submitted to chronic IVH demonstrated significantly elevated circulating levels of both IL-1β and IL-18 (Figure 5A, B), with similar concentrations to those observed after a single acute IVH event. Western blotting also identified an increase in NLRP3 protein in the livers of mice that underwent chronic IVH, compared to control mice (Figure 5C), but no significant alterations in other inflammasome components or precursor molecules were observed in these livers (pro-caspase-1, caspase-1 p20 or p15 unit, pro-IL-1β, IL-1β p15) (Supplementary Figure 1).

**Figure 5.**
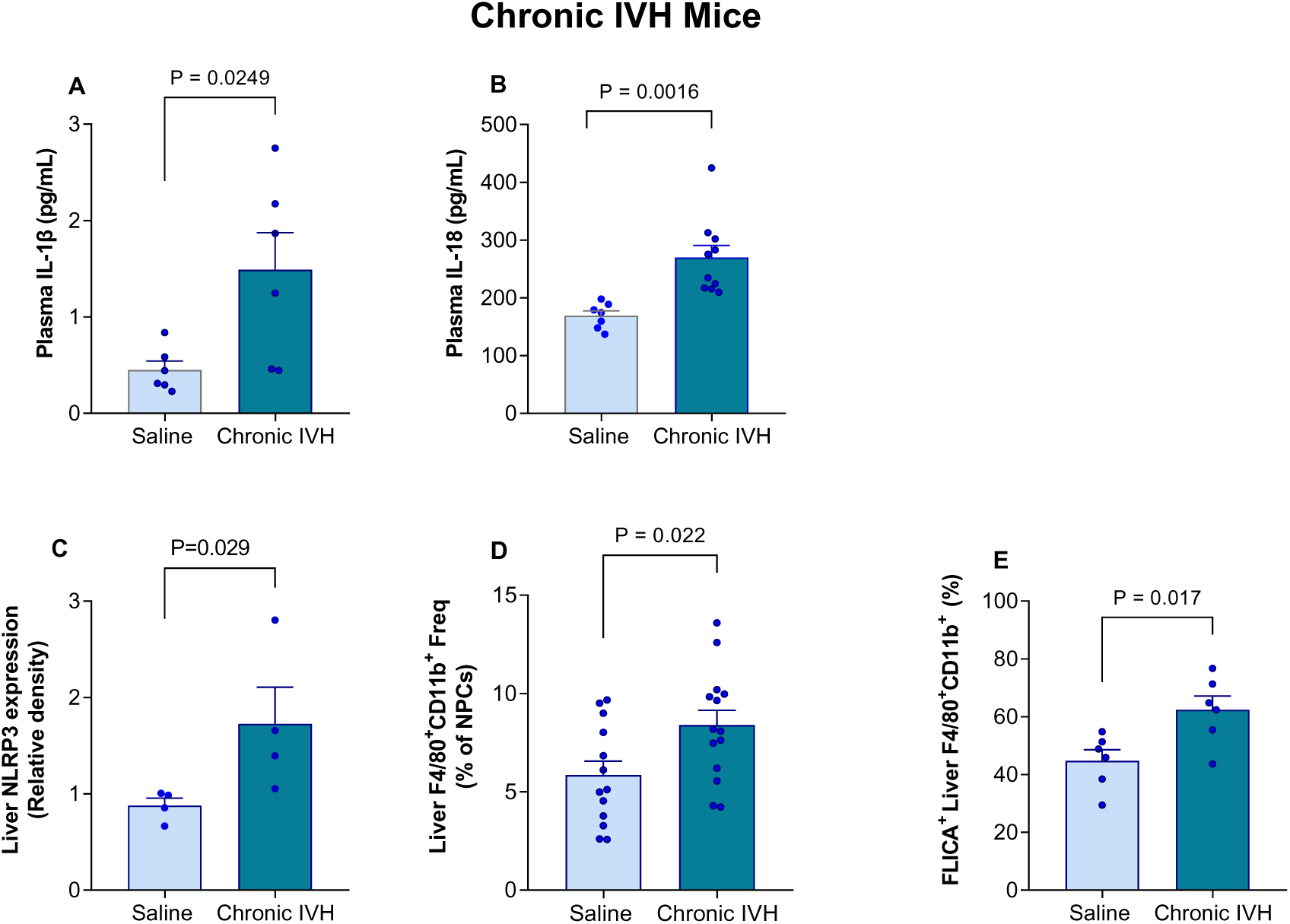
Chronic in vivo intravascular hemolysis induces liver and macrophage inflammasome activation. C57BL/6 mice were repeatedly administered (*i.p*.) with sterile 0.9% NaCl (CON, 2x/week) or low-dose phenylhydrazine (Chronic IVH, 10 mg/kg, 2x/week) for 4 weeks to induce chronic IVH, or not. Peripheral blood and liver tissue samples were separated from the mice at 48 hours after the last administration. Plasma IL-1β (A; n = 5-6) and IL-18 (B, n = 7-10) were quantified by ELISA. The protein expression of the NLRP3 inflammasome subunit was quantitated by Western blotting in the liver samples (C, n=4); expression was normalized by GAPDH expression in the same samples (Western blot membranes are displayed in Supplementary Figure 1). The frequency of F4/80^+^CD11b^+^ macrophages in non-parenchymal cell (NPC) suspensions obtained from the liver samples was determined by flow cytometry (D, n= 13-14). The percentage of these liver F4/80^+^CD11b^+^ macrophages displaying caspase-1 activity was determined using the FAM-YVAD-FLICA-FITC probe and flow cytometry (E, n=6).

Immunophenotyping by flow cytometry of murine liver demonstrated a significant increase in the number of infiltrated F4/80^+^CD11b^+^ cells in mice subjected to chronic hemolysis, compared to control mice. Furthermore, these F4/80^+^CD11b^+^ cells displayed a higher caspase activity in the hemolytic mice, as demonstrated by FLICA binding and flow cytometry (Figure 5D, E).

### Heme Induces Inflammatory Responses in Macro- and Microvascular Endothelial Cells

Considering the crucial roles of both leukocytes and the endothelium in vascular inflammation and leukocyte recruitment, we examined the inflammatory responses of human umbilical vein and microvascular endothelial cells (HUVEC and HMEC-1, respectively) to heme stimulation*, in vitro* (Figure 6). Following incubation with heme (50 μM hemin) for 3 hours, both HUVEC and HMEC-1 presented significant elevations in their surface expressions of the adhesion molecules ICAM-1 (Figure 6A and H), VCAM-1 (Figure 6B and I) and E-selectin (Figure 6C and J). These changes were associated with significant increases in the release of the inflammatory molecules, IL-1β, IL-6, IL-8 and MCP-1 from heme-activated HUVEC (Figure 6D-G), highlighting the inflammatory effect of the heme molecule on endothelial cells.

**Figure 6.**
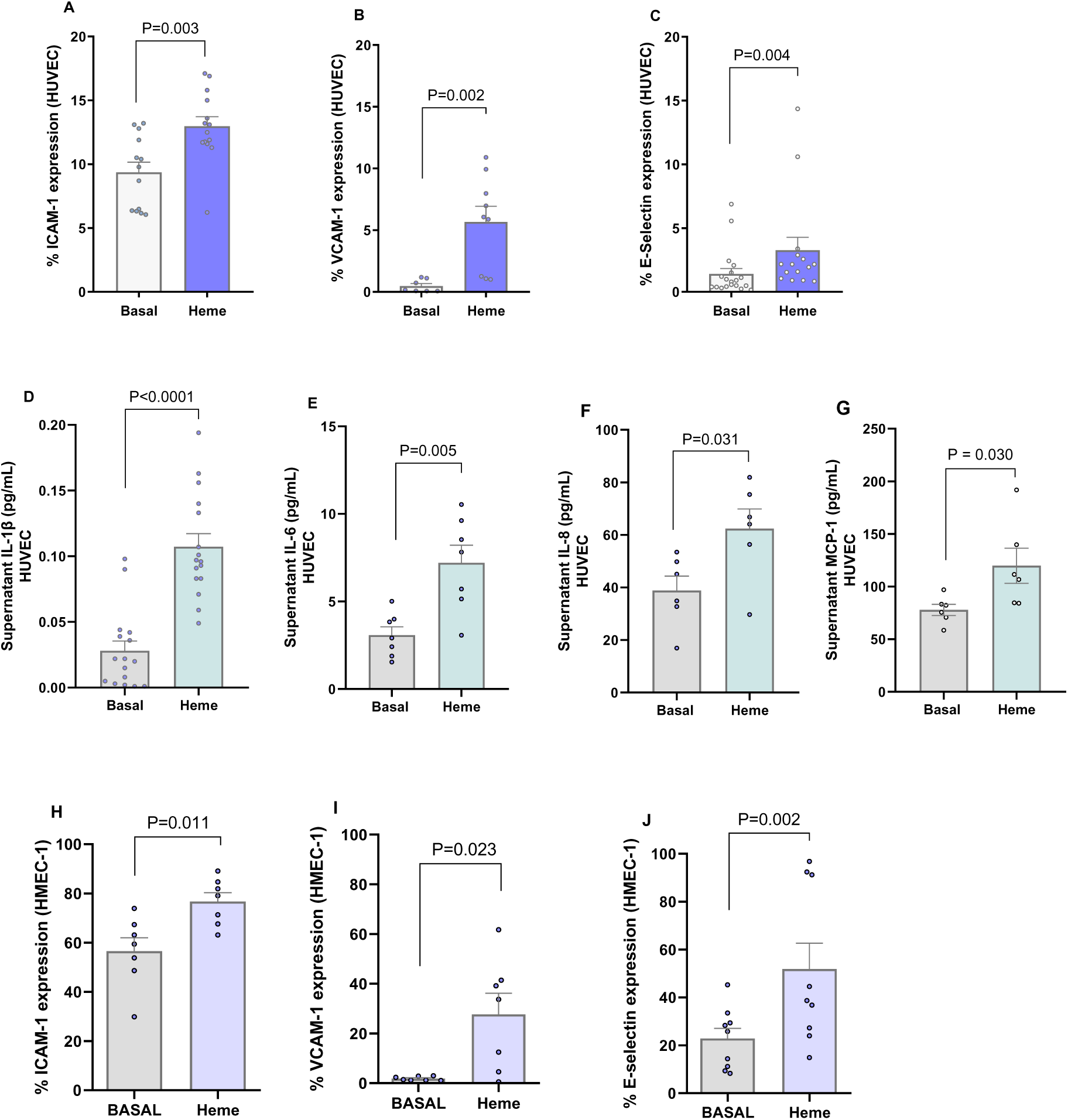
Effects of heme on endothelial cell activation, in vitro. Human umbilical vein endothelial cells (HUVECs, A-G) or human microvascular endothelial cells-1 (HMEC-1, H-J) were incubated under basal conditions (Basal), or with heme (50 μM) at 37°C, 5% CO₂ (3 hours). Cells (2 × 10⁵) were cultured in F12K medium supplemented with 2% fetal bovine serum (FBS). The percentage of cells expressing ICAM-1 (A, H), VCAM-1 (B, I), and E-selectin (C, J) was assessed by flow cytometry. Concentrations of inflammatory molecules, IL-1β (D), IL-6 (E), IL-8 (F), and MCP-1 (G) in HUVEC supernatants were determined by ELISA. Results are expressed as means ± SEM for duplicate or triplicate cultures and are representative of 6 to 15 independent experiments.

### Heme-Induced Adhesion Molecule Expression by Endothelial Cells is Dependent on Reactive Oxygen Species Generation and not Increased Caspase-1 Activity

To examine whether inflammasome activation participates in heme-induced inflammatory responses in endothelial cells, we determined caspase activity in heme-activated HUVEC and HMEC-1 using the FAM-FLICA probe and flow cytometry. Caspase activity was significantly increased in both HUVEC and HMEC-1 cells following heme incubation, compared to control cells (Figure 7A and 7H). Abrogation of heme-induced caspase activity in HUVEC by the YVAD inhibitor indicated a major role for caspase-1 in this caspase activity (Figure 7A). Importantly, qPCR revealed that the expression of the gene encoding caspase-1 in HUVEC was also elevated in response to heme stimulation (Figure 7B; gene expressions of other inflammasome components are shown in Supplementary Figure 2). However, inhibition of caspase-1 using the YVAD peptide, had no significant effect on the surface expressions of ICAM-1, VCAM-1 or E-selectin on heme-activated HUVEC (Figure 7E-G).

**Figure 7.**
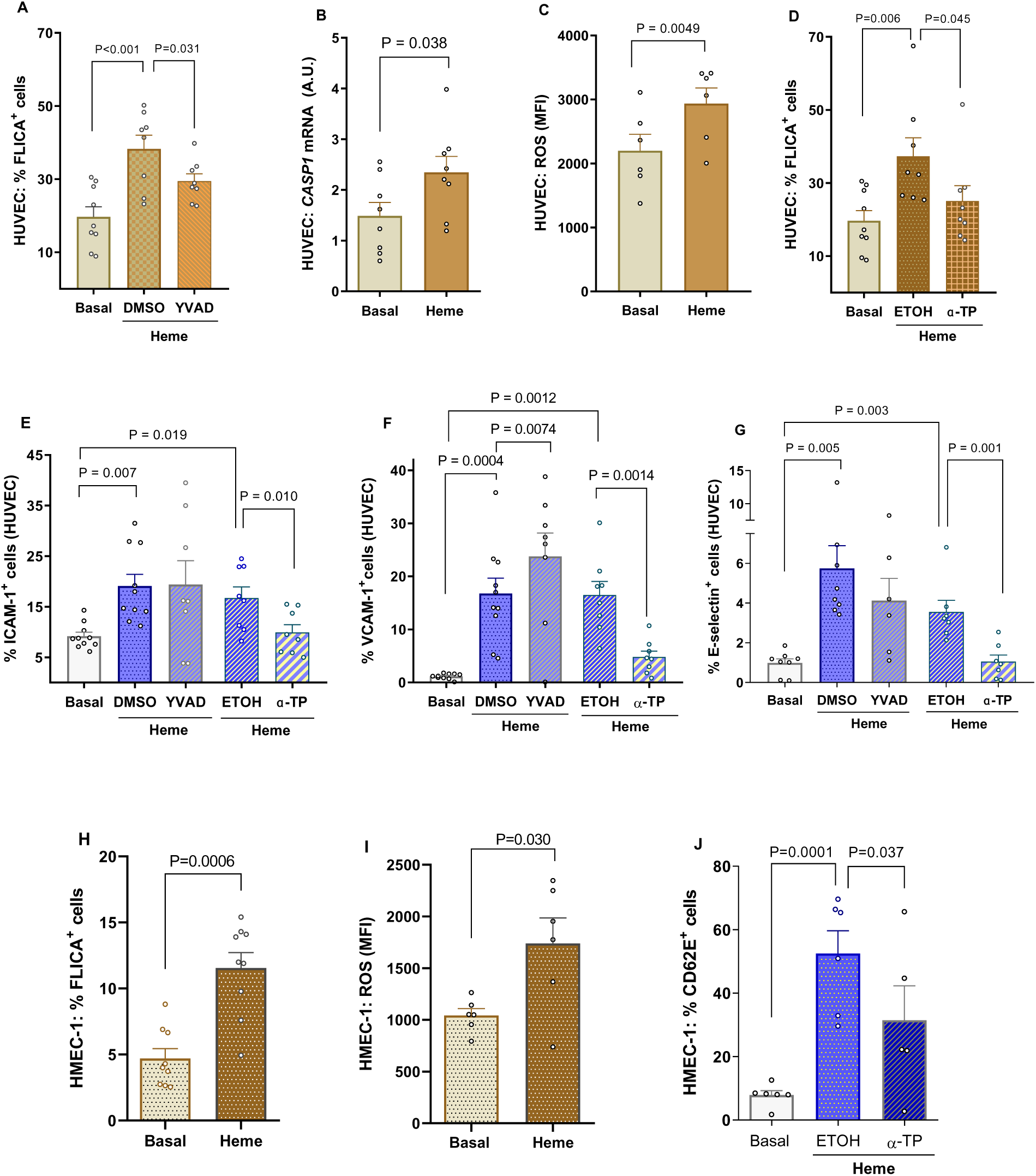
Role of Caspase-1 activation and ROS generation in heme-induced endothelial cell activation, in vitro. Human umbilical vein endothelial cells (HUVECs, A-G) or human microvascular endothelial cells-1 (HMEC-1, H-J) were incubated under basal conditions (Basal), or with heme (50 μM) at 37°C, 5% CO₂ (3 hours). Cells (2 × 10⁵) were cultured in F12K medium supplemented with 2% FBS. Cells were also pretreated for 1h with YVAD (50 μM), or DMSO (YVAD vehicle, 0.25%), or for 30min with α-tocopherol (α-TP, 1 mM) or ethanol (ETOH, α-TP vehicle, 2%) before stimulating with 50 μM heme. (A and H) Quantification of endothelial caspase-1 activity using the FAM-YVAD-FLICA-FITC probe and flow cytometry. (B) Quantification of the expression of the gene encoding caspase-1 in HUVEC by qPCR. (C and I) Quantification of reactive oxygen species (ROS) generation in cells using the DCFDA/H₂DCFDA probe and flow cytometry. Cellular expression of ICAM-1 (E), VCAM-1 (F), and E-selectin (G and J) was determined by flow cytometry.

Of note, the reactive oxygen species (ROS) content of both HUVEC and HMEC-1 were significantly amplified by heme activation (Figure 7C and I), as demonstrated using the DCFDA probe and flow cytometry. Importantly, neutralization of intracellular ROS generation by incubation with membrane-permeable α-tocopherol prevented the increase in caspase activity in HUVEC. In turn, in contrast to the effects of caspase-1 inhibition, the neutralization of ROS significantly abolished the inducing effects of heme on ICAM-1, VCAM-1, and E-selectin expression on HUVEC (Figure 7E-G) and abrogated heme-induced E-selectin expression on HMEC-1 (Figure 7J). Data suggest that the inflammatory effects of heme in endothelial cells may be, at least in part, mediated by the induction of oxidative stress and not necessary mediated by inflammasome/caspase activity.

## DISCUSSION

Although different hemolytic disorders have distinct clinical manifestations, some share certain characteristics, including inflammatory and pro-thrombotic processes, as well as manifestations such as pulmonary hypertension (1). Such manifestations are probably secondary to hemolysis, specifically IVH, itself irrespective of the cause and suggest that IVH acts an independent disease mechanism (43, 44). While the association between inflammation and hemolytic disorders, particularly sickle cell disease, is well documented (45–49), the specific role of the inflammasome in driving this inflammatory response is unclear (11, 29). Isolating IVH as a distinct disease mechanism provides a useful approach to study this process (17, 19). Using *in vivo* models of acute and chronic IVH, we characterized rapid and longer-term vascular inflammatory responses and explored a potential role for NLRP3-inflammasome activation in this mechanism.

As previously reported, our acute model of *in vivo* hemolysis induced moderate IVH in C57BL/6 mice, resulting in immediate elevations of plasma cell-free hemoglobin and total plasma cell-free heme (17). These levels were comparable to those of mice with sickle cell disease (17) and were accompanied by a rapid depletion of circulating haptoglobin, indicative of a mild acute hemolytic event. Circulating TNF-α, a potent pro-inflammatory cytokine (50), was up-regulated within 3 hours of IVH, suggestive of a systemic inflammatory response. Additionally, plasma S100A8, a leukocyte-derived putative biomarker for sepsis and a predictor of mortality in heart failure patients (51, 52), increased significantly within just 15 minutes of the acute hemolytic event, reflecting rapid leukocyte activation. Concomitantly, circulating IL-1β levels were augmented at 15 minutes post-IVH, indicative of increased inflammasome activity and cytokine processing.

Further supporting a role for an inflammasome-derived inflammatory response to hemolysis, liver IL-1β-cytokine content increased significantly within 1 hour in these mice, suggesting a potential role for hepatic macrophages in amplifying this inflammation (53, 54). Consistent with this hypothesis, mice subjected to continuous IVH for 4 weeks exhibited an increased frequency of hepatic F4/80^+^CD11b^+^ cells and a subset of these cells displayed augmented caspase activity; indeed, NLRP3-inflammasome activation mediates heme-induced macrophage activation (55). Interestingly, although IL-18 also increased post hemolysis in mice, it was only detectable in the circulation only at 6 hours after the event. This delay may reflect the need for gene expression upregulation to mediate increased pro-IL-18 production (56). However, continuous IVH for 4 weeks triggered a persistent rise in plasma IL-1β and IL-18 levels, accompanied by the upregulation of NLRP3 gene expression in the liver, highlighting a continuous role for inflammasome activation during hemolytic processes.

Given that the rapid alterations observed suggested leukocyte participation in the hemolytic inflammatory response, and considering the concomitant upregulation of the S100A8 alarmin, which may prime the inflammasome (57), we monitored caspase activity in the neutrophils and monocytes of the mice following induction of acute hemolysis. Caspase-1 is recruited and activated by inflammasome complexes in response to PAMPs or DAMPs, such as the heme molecule. Caspase activity, measured using FAM-FLICA YVAD-FMK-probe binding (58), was significantly elevated in both leukocyte types at one hour after IVH. Furthermore, both the caspase-1 inhibitor, YVAD, and the NLRP3 inhibitor, MCC950, completely abolished the effect of hemolysis on FLICA probe binding in both neutrophils and monocytes of mice following acute IVH, suggesting that leukocyte NLRP3-inflammasome assembly is triggered by this event and may contribute to the observed elevation in circulating IL-1β cytokine. It should be noted that while the caspase-1 inhibitor, AC-YVAD-CMK, may exhibit cross reactivity with other caspases (-4, -5 in humans)(59), it is unlikely that such off target effects occurred at the administration dose employed.

The NLRP3 inflammasome has been implicated in leukocyte recruitment and migration mechanisms in response to inflammatory triggers (60, 61) and previous studies suggest that heme can induce neutrophil adhesive mechanisms (23). Using intravascular microscopy techniques to assess the microvascular circulation in the cremaster muscle of mice, we confirmed the immediate effect of acute IVH on microvascular leukocyte recruitment (17). To assess the role of NLRP3-inflammasome-mediated activity in this event, we inhibited caspase-1 or NLRP3 activity before inducing IVH. Importantly, caspase-1 inhibition significantly abrogated hemolysis-induced leukocyte rolling and adhesion at the vascular wall, whereas NLRP3 inhibition had a more limited effect, only partially reducing hemolysis-induced leukocyte recruitment in the microvasculature. While we have previously reported the rapid effect of IVH on microcirculatory leukocyte recruitment in mice (17), its impact on microcirculatory blood flow and tissue perfusion has not been reported. Using laser Doppler flowmetry to monitor blood flow dynamics, we show for the first time that acute IVH leads to significant and rapid reductions in blood flow velocity and blood perfusion within the cutaneous microcirculation of mice. These findings suggest the onset of an acute tissue ischemic processes, potentially mediated by the observed vascular leukocyte recruitment to the endothelium (64, 65). A role for NO consumption in this rapid hypoperfusion is plausible, given the established effect of hemolysis and cell-free Hb release on vascular NO levels (66, 67). Indeed, 6 hour-water infusion in dogs significantly elevates mean arterial pressure, and this effect can be attenuated by NO inhalation (3); however, both the YVAD caspase inhibitor and the MCC950 NLRP3 inhibitor restored cutaneous blood perfusion in mice subjected to IVH. Thus, while the caspase-dependent microvascular leukocyte recruitment induced by IVH may not be entirely dependent on the NLRP3 inflammasome, NLRP3-inflammasome-mediated inflammatory responses played a significant role in the resulting impairment of skin microvascular hemodynamics observed following IVH. These differences may reflect distinct molecular mechanisms and temporal dynamics governing leukocyte recruitment and migration, and ischemic processes. Importantly, impaired skin perfusion as a result of IVH may contribute to cutaneous lesions, such as skin ulcers (68, 69), a feature of some diseases that display IVH (70, 71).

Having identified roles for NLRP3-inflammasome and caspase-1 activation in IVH-induced microvascular leukocyte recruitment and ischemia *in vivo*, potentially involving endothelial cell interactions, we sought to explore the effects of heme on endothelial cell activation and inflammasome activity, *in vitro*. Endothelial cells are reported to display NLRP3 inflammasome assembly under certain inflammatory conditions (72, 73), and this activity may contribute to their adhesive interactions. The incubation of both macrovascular (HUVEC) and microvascular (HMEC-1) endothelial cells with heme at a concentration similar to that found in the circulation of patients with sickle cell disease and mice induced to acute hemolysis prompted significant upregulation of the surface expressions of the adhesion molecules, ICAM-1, VCAM-1 and E-selection, which are all important for mediating leukocyte adhesive interactions with the vascular endothelium (74). Furthermore, incubation with the heme molecule stimulated the production of inflammatory chemokines and cytokines by endothelial cells, reaffirming its role in endothelial cell activation (14).

Notably, heme upregulated caspase-1 gene expression in macrovascular endothelial cells and augmented caspase activity in both the HUVEC and HMEC-1 cells. This was accompanied by the generation of ROS, confirming a role for oxidative stress in mediating heme-induced endothelial activation (14, 36). Importantly, pre-incubation of HUVEC with the anti-oxidant, α-tocopherol, reduced caspase activity in a similar manner to the inhibition of caspase-1, indicating that, in fact, the generation of ROS in response to heme may be playing an intracellular signaling role in endothelial caspase-1 activation, consistent with the recognized role of ROS-triggered NLRP3 inflammasome activation in endothelial dysfunction (34). Additionally, while caspase-1 inhibition by YVAD had no significant effect on heme-induced vascular endothelial cell adhesion molecule expression, a membrane-permeable antioxidant significantly reduced the surface upregulation of ICAM-1, VCAM-1 and E-selectin on these HUVEC cells and partially abrogated the upregulation of E-selectin on HMEC-1. As such, while ROS-induced caspase-1 activity is associated with endothelial cell activation by heme, the effect of this DAMP on adhesion molecule expression appears to be mediated directly by ROS generation and signaling rather than by an inflammasome/caspase-1–dependent pathway. A role for NF-κB pathway signaling has been demonstrated in heme-induced neutrophil adhesion molecule expression (23), and has also been implicated in ROS-induced endothelial cell activation in response to other molecular patterns such as oxidized low density lipoprotein (oxLDL) (75) and LPS (76).

Our findings suggest that caspase-1 activation and rapid IL-1β processing play a significant role in mediating inflammatory responses to IVH, contributing to key outcomes such as reduced tissue blood perfusion and microvascular leukocyte recruitment. Moreover, NLRP3-inflammasome activation appears to drive this caspase-1-dependent leukocyte activation, particularly in the context of hypoperfusion. While ROS generation appears to primarily regulate vascular endothelial adhesion molecule expression in response to heme, caspase-1 activation may contribute to broader inflammatory consequences in these cells. Given the significance of IVH in driving inflammatory responses across different disorders, further studies are warranted to improve therapeutic strategies that reduce IVH and mitigate the effects of RBC DAMP release. In addition to complement therapy (77), in use for certain hemolytic diseases, promising approaches include hemopexin therapy, currently in clinical trials [NCT06699849]. Targeting inflammasome- and caspase-dependent pathways may further help attenuate the downstream effects of IVH, with potential for maintaining endothelial integrity, limiting leukocyte activation, and reducing the ischemic injury associated with IVH.

## DATA AVAILABILITY

Datasets will be made available from the corresponding author upon reasonable request.

## SUPPLEMENTAL MATERIAL

Supplemental Data is provided in the “Supplemental Material” section.

## ACKNOWLEDGMENTS

The authors would like to thank Irene Santos, for assistance with flow cytometry analysis.

## AUTHOR CONTRIBUTIONS

PLB and NC conceived and designed the research. PLB, LFSG, VF, FCL, and EMFG performed experiments. PLB and NC analyzed data. PLB, FFC, and NC interpreted the results of the experiments. PLB and NC drafted the manuscript. All authors edited and revised the manuscript and approved the final version.

## Supplementary Material

**Supplementary Table 1.** List of antibodies for flow cytometry.

| Antibodies | Source | Catalog |
| --- | --- | --- |
| Rat Monoclonal anti-mouse/human CD11b (M1/70), APC | BD Pharmingen | Cat# 553312; RRID: AB_398535 |
| Rat Monoclonal Anti-mouse Ly-6G Antibody, PE | BioLegend | Cat# 127608; RRID: AB_1186099 |
| Rat Monoclonal CD43 Monoclonal Antibody (eBioR2/60), PE | eBioscience | Cat #12-0431-82; RRID: AB_465659 |
| Rat Monoclonal Ly-6C Monoclonal Antibody (HK1.4), APC | eBioscience | Cat# 17-5932-82; RRID: AB_1724153 |
| Rat Monoclonal F4/80 Monoclonal Antibody (BM8), PE | Invitrogen | Cat# 12-4801-82; RRID: AB_465923 |
| Rat Monoclonal Anti-mouse/human CD11b Antibody, PerCP/Cyanine5.5 | BioLegend | Cat# 101228; RRID: AB_893232 |
| Mouse anti-Human CD62E (E-Selectin), PE | BD Pharmingen | Cat# 551145; RRID: AB_394072 |
| Mouse anti-human ICAM-1 Monoclonal Antibody (1H4), PerCP | Invitrogen | Cat# MA1-10250; RRID: AB_11156372 |
| Rat Mouse Anti-Human CD106 (51-10C9), APC | BD Pharmingen | Cat# 551147; RRID: AB_398496 |

**Supplementary Table 2.**
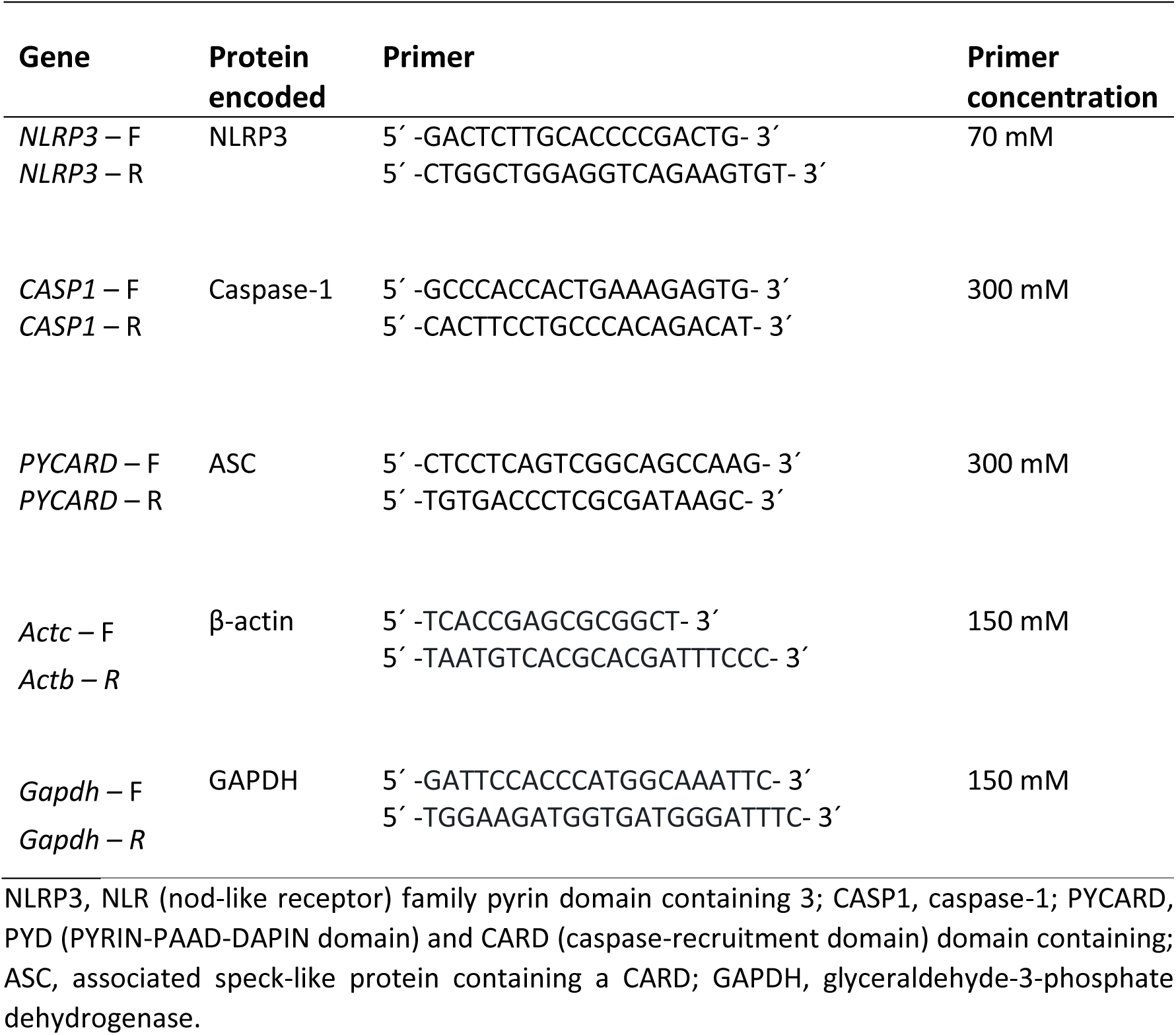
Primer sequences for quantitative Real-Time PCR.

**Supplementary Figure 1.**
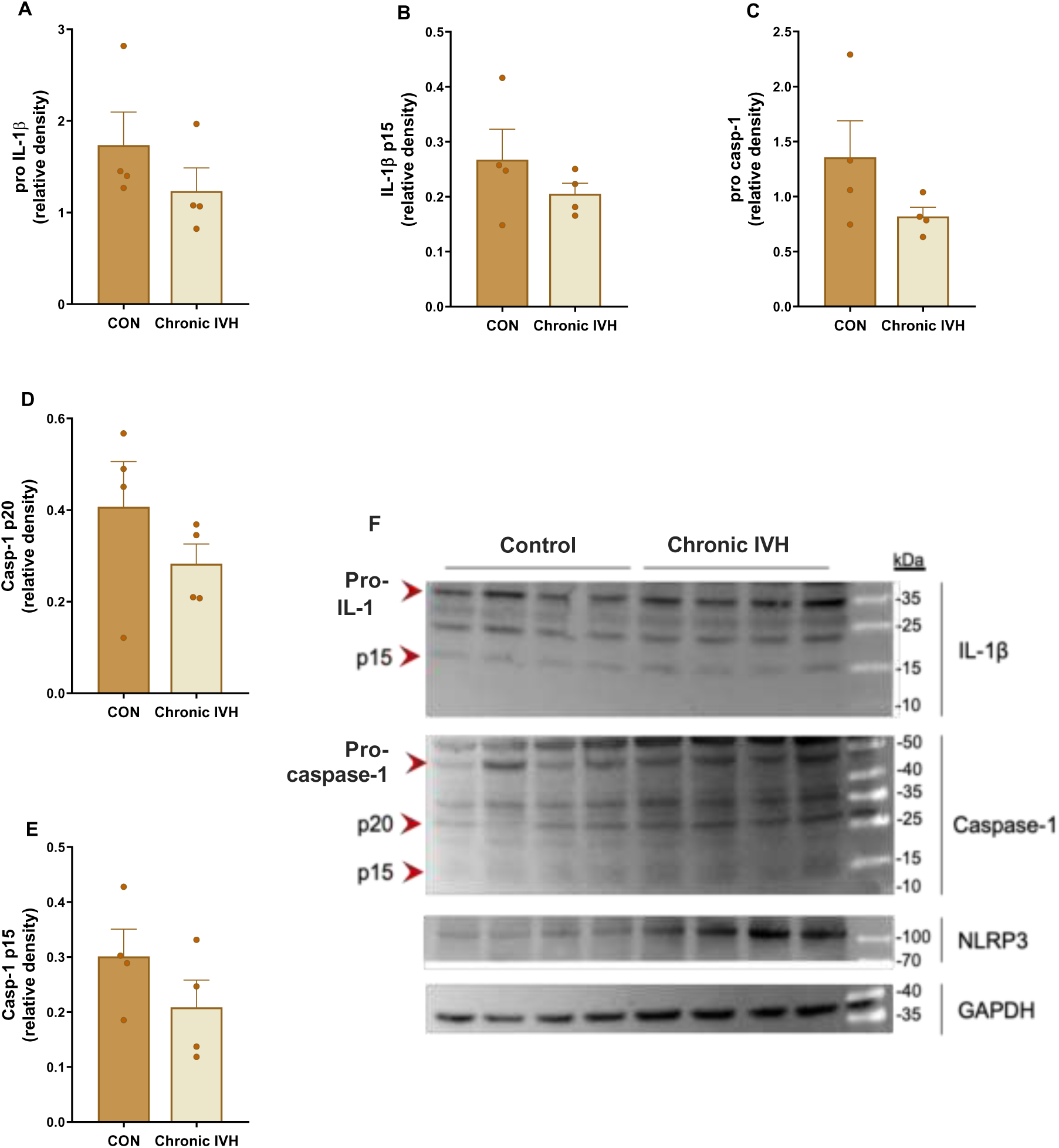
Effect of chronic *in vivo* intravascular hemolysis on liver inflammasome components. C57BL/6 mice were administered (*i.p*.) with sterile 0.9% NaCl (CON, 2x/week) or low-dose phenylhydrazine (Chronic IVH, 10 mg/kg, 2x/week) for 4 weeks to induce chronic IVH, or not. Liver samples were separated from the mice at 48 hours after the last administration, and inflammasome component protein expressions were quantitated by Western blotting (Control, n=4; Chronic IVH, n=4); expression was normalized by GAPDH expression in the same samples. (A) pro-IL-1β, (B) cleaved IL-1β (p15 subunit), (C) pro-caspase-1, and the active forms; (D) caspase-1 p20 subunit, (E) caspase-1 p15 subunit. (F) Western blot membranes incubated with primary antibodies for the respective proteins.

**Supplementary Figure 2.**
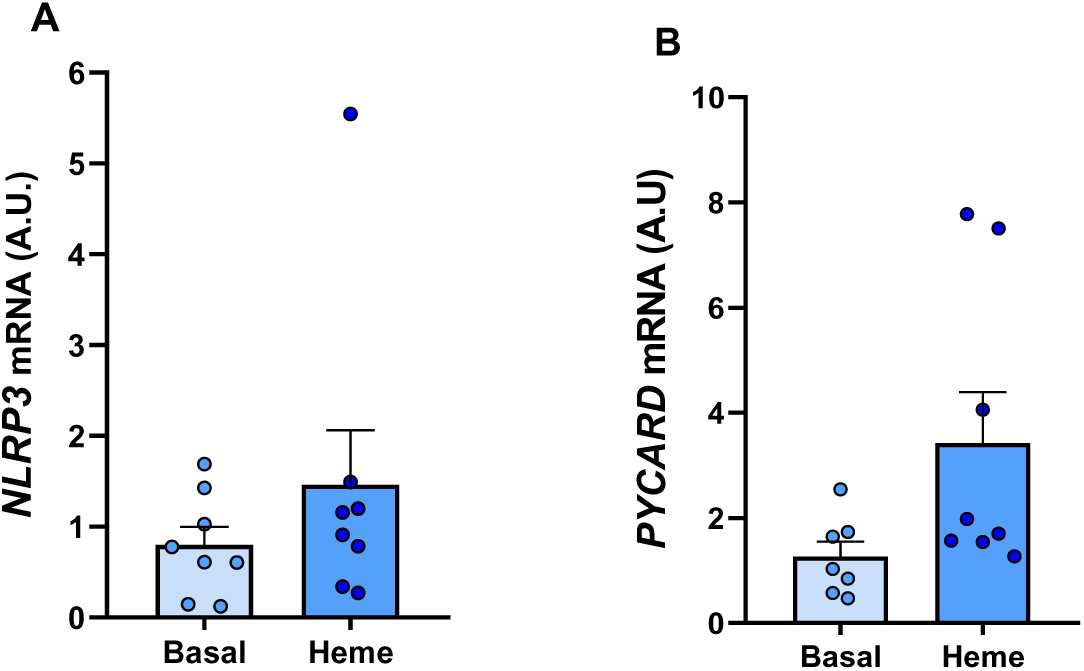
Effect of heme on the induction of gene expressions of inflammasome components in human umbilical vein endothelial cells (HUVEC). Human umbilical vein endothelial cells were incubated under basal conditions (Basal), or with heme (50 μM) at 37°C, 5% CO₂ (3 hours). Cells (2 × 10⁵) were cultured in F12K medium supplemented with 2% FBS. The mRNA expressions of (A) *NLRP3* (n=8) and (*B) PYCARD* (n=7-8) were determined by quantitative PCR and normalized using the housekeeping genes β-actin and GAPDH, with gNorm.

**Supplementary Videos. Links for representative videos of leukocyte recruitment in the cremaster muscle microcirculation of C57BL/6 mice after procedures.** C57BL/6 mice were pretreated (*i.p*., 1 hour) with YVAD (8 mg/kg) or MCC950 (50 mg/kg) or with DMSO vehicle (0.5% in sterile PBS, vehicle for YVAD), or with sterile PBS vehicle. Intravital microscopy was performed on the cremaster muscle after intravenous administration of sterile 0.9% NaCl (CON) or after the hemolytic stimulus (IVH).

**Video 1: CONTROL + DMSO.** Representative venule from a mouse that was pretreated with DMSO and then received i.v. sterile 0.9% NaCl (at approximately 15-20 min after i.v. administration). https://drive.google.com/file/d/1XMVaxZOryRYkxfMHwZ5NQ65MLA4jWg05/view?usp=drive_link

**Video 2: Acute IVH + DMSO.** Representative venule from a mouse that was pretreated with DMSO and then received i.v. sterile water to induce intravascular hemolysis (at approximately 15-20 min after i.v. administration). https://drive.google.com/file/d/1Cq6xHoXnNXD6_Fr--_Cm6PHXD-EBYcow/view?usp=drive_link

**Video 3: Acute IVH + YVAD.** Representative venule from a mouse that was pretreated with YVAD and then received i.v. sterile water to induce intravascular hemolysis (at approximately 15-20 min after i.v. administration). https://drive.google.com/file/d/1i-KpHZD8gopEXWuXhsP-LZMetfDF_15d/view?usp=drive_link

**Video 4: CONTROL + PBS.** Representative venule from a mouse that was pretreated with PBS and then received i.v. sterile 0.9% NaCl (at approximately 15-20 min after i.v. administration). https://drive.google.com/file/d/1GbBxTx_U_zqUe8vtRt2xPUHpxRBvVwwb/view?usp=drive_link

**Video 5: Acute IVH + PBS.** Representative venule from a mouse that was pretreated with DMSO and then received i.v. sterile water to induce intravascular hemolysis (at approximately 15-20 min after i.v. administration). https://drive.google.com/file/d/1Iot9KE-Gvu8sDC885k12XYvN-FbP1Rfa/view?usp=drive_link

**Video 6: Acute IVH + MCC950.** Representative venule from a mouse that was pretreated with MCC950 and then received i.v. sterile water to induce intravascular hemolysis (at approximately 15-20 min after i.v. administration). https://drive.google.com/file/d/1Mv0wPR7dpnZ8ooOuurCHyTKZzbF_QUtQ/view?usp=drive_link

